# Enabling Fast Annotation Process With Table2Annotation Tool

**DOI:** 10.1101/2020.04.03.023069

**Authors:** Pierre Larmande, Kazim Muhammed Jibril

## Abstract

Semantic annotation is the process in which semantic concepts are linked to natural language. It helps in boosting the search and access of resources and can be used in information retrieval systems to increase the queries from the user. In this paper, we are interested in identifying ontological concepts in scientific text contained in spreadsheet. We developed a tool which is able to handle various types of spreadsheet. Furthermore, we used the benefits of NCBO Annotator API provided by BioPortal to enhance the semantic annotation functionalities covering spreadsheet data. Table2Annotation developed strengths in certain criteria like speed, error handling and complex concept matching.

**Availability:** GitHub : https://github.com/pierrelarmande/ontology-project

## Introduction

Semantic annotation have various definitions from various authors in various papers but then they all similar with one clear purpose. To define this, from [1], semantic annotation is the process in which semantic concepts are linked to natural language. [2] defines semantic annotation as methods of describing resources (texts, images, etc.) with meta-data where the meaning has been specified in an ontology. According to [1] semantic annotation can be seen as a methodology of inputting meta-data which are classes, properties, relations and instances (i.e concepts of an ontology) in web resources to be able to give or allocate semantics. Summarizing all definitions we can simply say that semantic annotation is a way of matching resources to ontologies.

To make it more clearer take this example of text *“*..*days to flowering*..*”*, with the help of semantic annotation we would be able to match this text to the ontology concept id from the Trait Ontology [3]*”TO:0000344”* which indicates the concept *“days to flowering trait”*.

Semantic annotation helps in boosting the search and access of resources. From [4], Semantic annotation can be used in information retrieval systems to increase the queries from the user with some ontology terms and also provide grouping of documents gotten based on specific contents. In Biomedical resources like there are a lot of abbreviations in the texts which makes it hard for researches to comprehend. In these texts you can find an abbreviation with a lot of meanings not just one. Semantic annotation helps us to be able to make clear and understand abbreviated terms based on the way they appear in a context.

In this paper, we want to be able to identify ontological concepts in scientific text, it could be seen as an ontology matching process to match texts with concepts and there are already some existing web services and tools that uses semantic annotation for ontology matching which are been evaluated in [1].

The paper is organized as follows. Section 2defines the challenges of semantic annotation. Section 3 presents an overview of Table2Annotation. Section 4 analyses the results of a semantic annotation example. Section 5 concludes the manuscript.

## Semantic Annotation Challenges

There are benefits of semantic annotation but there are also some challenges faced during annotation of biological texts or other resources. Some of these challenges are:

- Text ambiguity in bio terms. Terms usually has more than one meaning causing ambiguity making it unclear and difficult to annotate.
- Given texts having incorrect spellings or grammar.
- Positions of words in a sentence for example *“drought and salinity tolerance”* means *“drought tolerance and salinity tolerance”* but in this case we might have matching on only *“salinity tolerance”*.
- In biomedical context, all protein has associated genes with the same name making it difficult for annotation on gene and protein texts.
- Abbreviations and short hand texts can also complicate annotations.

The challenges faced in annotation can be tackled by two methods which are term-to-concept matching method which has to do with matching some parts of provided texts to structured knowledge databases, dictionaries or vocabularies and machine learning (ML) method which has to do with creating annotators for specific purposes and usage instead of general usage [4].

Specifically for the 3rd challenge, it can be tackled by creating algorithms that can transform texts with conjunctions like *“and”* or *“or”* so in the example of *“drought and salinity tolerance”*, this algorithm should be able to transform this phrase to *“drought tolerance and salinity tolerance”* before annotation proceeds.

Although these are good solutions to tackle some challenges but also there are some drawbacks too. Drawbacks of the term-to-concept matching is its inability to disambiguate terms so annotators that inherit this method usually match terms with several possibilities. This drawback is encountered in the use of the NCBO annotator [5] and one way to solve this problem is to be able to have several algorithms that use some knowledge based dictionaries to transform ambiguous terms to similar meaning making it clear for the annotator. This algorithms should also be able to correct incorrect grammars and wrong spellings by matching dictionary terms with similar spellings or phrases.

### Challenges of Semantic Annotation Tools

There are diverse tools used in semantic annotation [1]. These tools also have some challenges faced when using them and to list a few as follows:

- **Speed**. This is one of the most common challenge. Annotations performed on huge datasets can take a lot of time to finish.
- **Language specific**. Most of the annotators are in English which makes applying semantic annotation in other languages a challenge.
- **Document format**. Annotators that support input of documents can face the problem of having to annotate different document formats and not supporting a particular format could be a challenge.
- **Text Variation**. According to [4], challenges are faced also due to the fact that there are different kinds of biomedical texts and variations among variations of text for example in biomedical and clinical text.
- **Disambiguation**. Entities mentioned in biomedical texts sometimes don’t have enough context for under-standing.

These challenges and more others are been studied and so many experts try to figure out the way to tackle them in new systems developed. These challenges may not be fully tackled but can be reduced and the following section shows what we did to tackle some of the challenges in the developed system in this project.

## Overview of Table2Annotation tool

In this section, we describe the solution proposed to build the ontology matching system. Our solution is using NCBO annotator Web Service API as a primary information retrieval.

NCBO annotator annotates data with MGrep term-to-concept matching tool and retrieves sets of annotations which are later expanded using various methods of semantic matching making this annotator to go through 2 stages. This is a unique annotator because of the method used to associate concepts instead of looking for a concept which will best match the provided context. This annotator uses BioPortal [6] and UMLS Metathesaurus ontologies vocabularies and although it doesn’t support disambiguation of terms, it is suitable for real time processing. This annotator is available for free and is implemented through web services. This annotator can be used on AgroPortal [7] and BioPortal.

The overview diagram of the Table2Annotation tool is shown on supplementary file S1. The flow of the system is quite simple and understandable. The system starts by taking an input dataset which is a file in CSV, Excel, etc. and then processes this file by reading the data and fetching the necessary data to be annotated. It takes the necessary data and calls an external API provided by AgroPortal to annotate this data. The results returned from this process is processed by taking the URI, concept-id and the matched words. Finally the annotated terms are being saved and written to an output file for the user to access.

The operation of the matching system is described diagrammatically on supplementary file S2. In building this Table2Annotation tool we decided to use the NCBO annotator (AgroPortal API) to support annotation of terms.

### Important Algorithms

As seen in the challenge section, there are several problems which are to be handled and to handle this problems some special algorithms where created to handle some of them.

#### Threading

First of all the system was created in functional independent approach where major functions are independent like getting inputs and annotation are independent. This allows us to be able to handle the part of the system which is slow. The part or function which slows down the systems is that function that deals with iterating through the cells, taking the cell data and then annotating this data. To be able to reduce the problem of speed (Problem 1) we decided to create an algorithm to speed up the process. The algorithm uses Multi-Threading concept allowing the function to be run by several processors (threads) concurrently.

#### Permutation

As seen before the problem of grammar (Problem 2), although the acronym and plural problems haven’t been solved yet and been reserved for future enhancements the problem of conjunctions can be reduced by creating an algorithm to handle this case.

#### Multiple Dataset Formats

The problem of dataset formats (Problem 5) was reduced by an algorithm to detect what file format is being in- putted by the user and then handling the process depending on the file format.

### Running Table2Annotation

Table2Annotation is a Java based program which is currently executed through command line interface. The user needs to have a dataset which he/she wants to annotate first before anything. Table2Annotation is compiled after the code and all the functions explained in previous chapter have been fully implemented. The compilation of Table2Annotation is done with all the necessary libraries included in the java project. To run the system the user needs to input as parameters the following; *Input file (mandatory), Column (mandatory), Suggestions (optional), Slice (optional), Separator (optional), Sheet (optional)*.

Firstly the user provides the path to the input file (dataset) and then secondly provides the name of the column to be annotated. These two parameters are mandatory and the rest are optional. If the user wishes to explore the other functions the user can pass as parameters the suggestions (recommendations) ontologies, the slice (grouping), the separator if the file is a separated file type and the sheet number if its an excel file with multiple sheets. After the command is executed the system starts processing and stops when the process is completed. The results of this operation is outputted in a file in same format as the input file and given to the user.

## Results

In this section we describe the results gotten from our Table2Annotation tool. It also describes the context to obtaining the results and an evaluation of the system.

First of all to begin test running the system, we need to have a dataset to test with and this is shown in figure 1. The data is quite small as we want to use a small data to better show clarity on the results gotten but its still the same using a large dataset anyways. The dataset contains a *“PROPERTY”* column which is the column with the terms that will be annotated.

**Figure 1:**
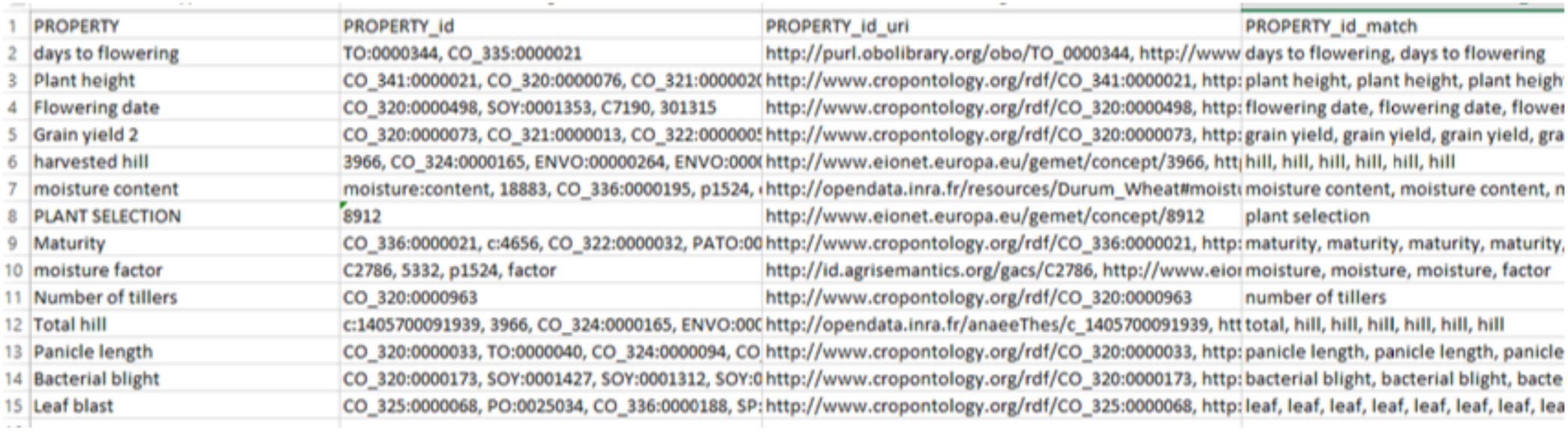
Dataset Without Suggestion and Slice.

### Test Without Recommendation or Slice

In this test, we ran the system without giving recommendations or slice options (i.e ontology list to map on provided by AgroPortal) and the results of this test can be seen in figure 1. In the results of the operation we can see that there are 3 new columns which are the *“PROPERTY_id”, “PROPERTY_id_url”* and *“PROPERTY_id_match”*. The first added column contains the concept-ids gotten from the annotation, the second added column contains the uri’s of the concepts and the third added column contains the matched terms with the concept.

### Test With Slice

In this test we test ran the system by giving a slice called *“agrold”* which is a slice containing ontology groups for agronomy. The results of the test is shown in figure 2. In the results we can see three terms (highlighted in yellow) having no matching with any concept and this is because they do not have ontologies belonging to the *“agrold”* group.

**Figure 2:**
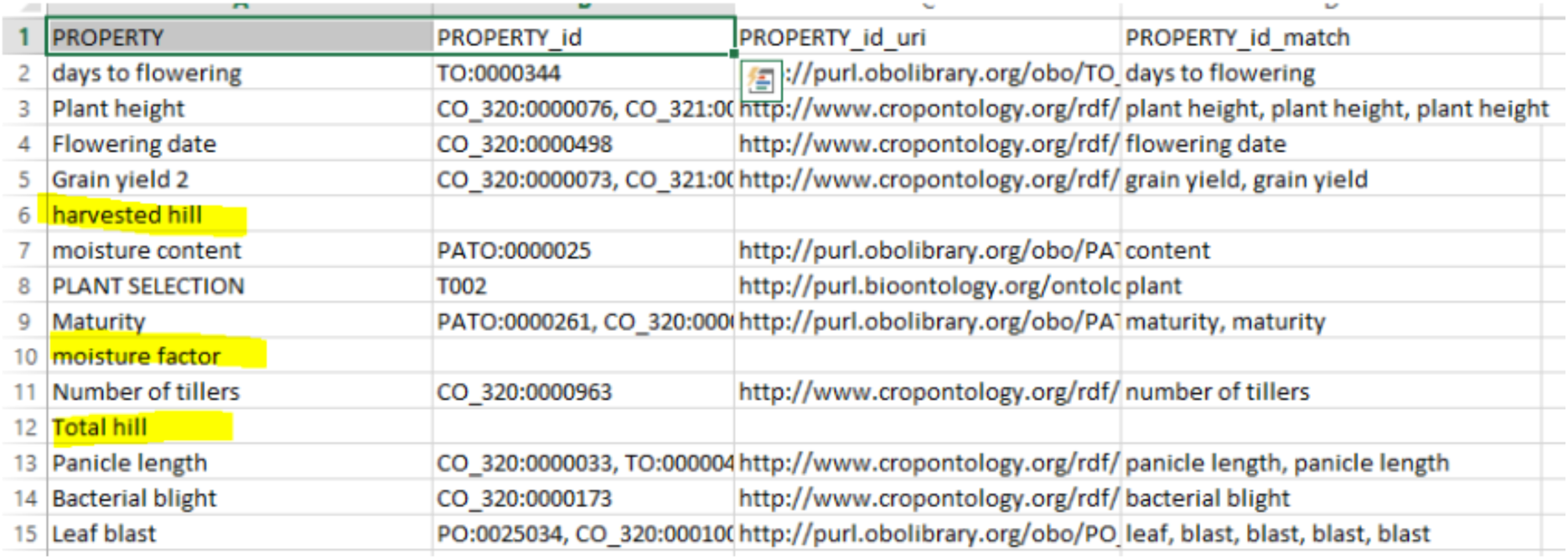
Dataset Result With Slice.

### Test With Recommendation

In this test we test ran the system by giving recommendations or suggestions. We tested by giving three suggestions which are *“PO (Plant Ontology)”, “TO (Plant Trait Ontology)”* and *“PATO (Phenotypic Quality Ontology)”*. The results of the test is shown in figure 3. In the results we can see that 6 terms (highlighted in pink) has no matching concepts and this is because we filtered the annotation to the three ontologies giving in the suggestions.

**Figure 3:**
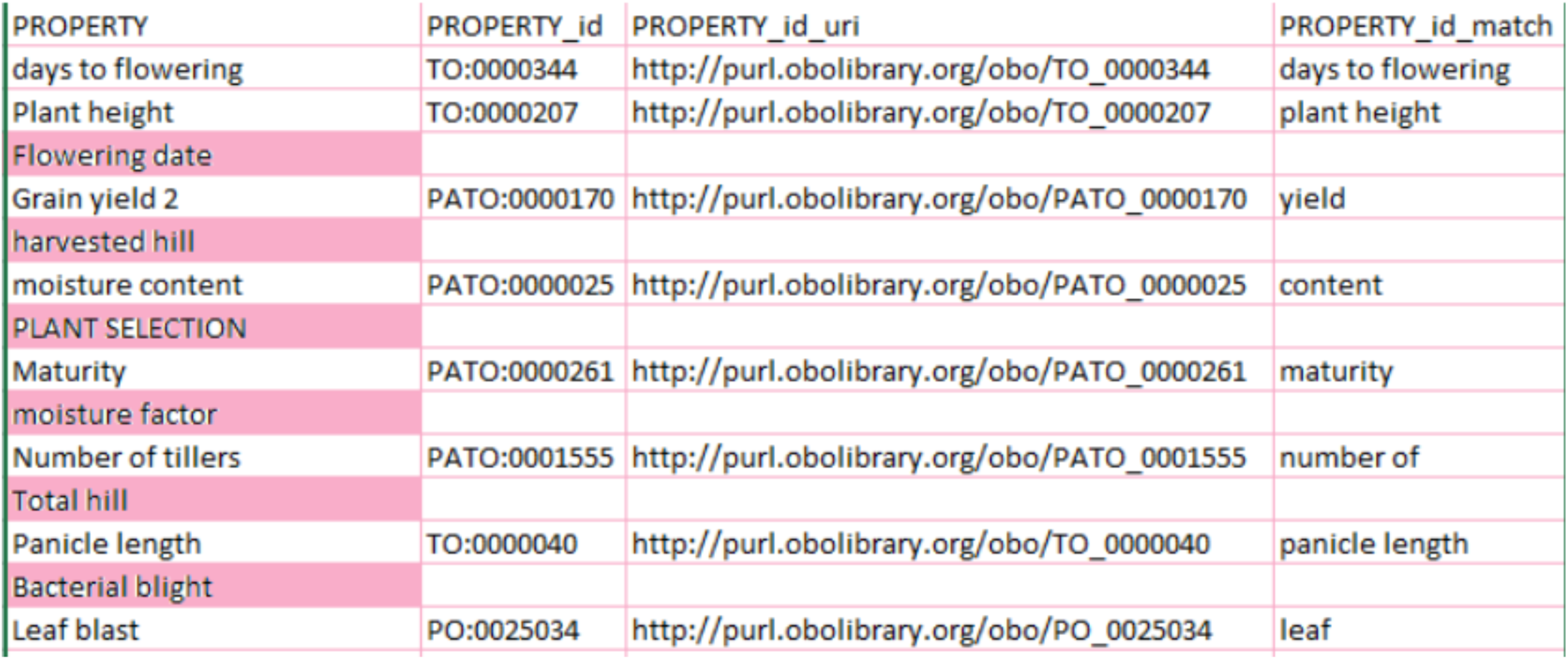
Dataset Result With Recommendation.

### Test With Permutation Algorithm

In this section we try to show the effect of having an algorithm to solve the problem of conjunction in terms which was mentioned in earlier (Problem 2). We annotate the term *“drought and salinity tolerance”* and figure 4 shows the results. Figure 4B shows the results from the operation with the algorithm and we can see that we have matching for three (3) terms which are *“drought”, “salinity tolerance”* and *“drought tolerance”*. Figure 4A shows the results from the operation without the algorithm and we can see that we have matching for just two (2) terms which are *“drought”* and *“salinity tolerance”*.

**Figure 4:**
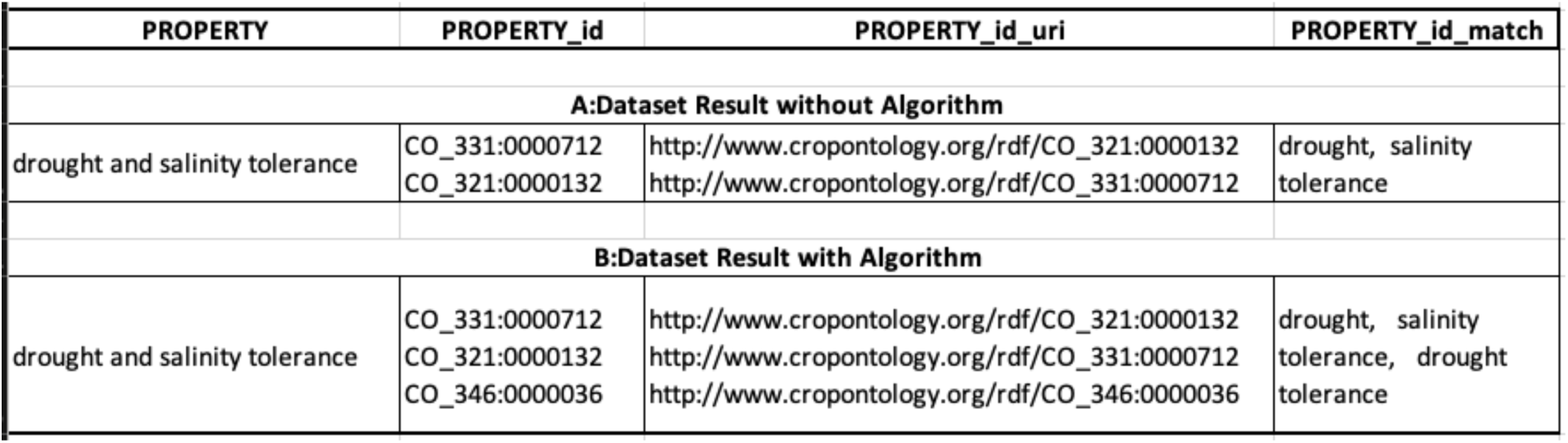
Dataset Result with and without Algorithm.

## Conclusion

In conclusion, Table2Annotation has strengths in certain criteria like speed, error handling and concept matching. Firstly as known already we use multi-threading algorithm making it to run process very effectively and efficiently. Secondly it handles errors and exceptions by ignoring them whenever there is one. If there is an error while matching one term it skips the term with error and continues to the next one while if there is a general error it still completes matching process but returns empty results. This way of error handling helps the user to run the process and do other things and come back to take the results not having to worry about system process terminations. Lastly the matching results are good and we see that cases of conjunctions are handled making the result contain more matches. The filters (slice and suggestions) also help to filter the results according to users expectations.

The systems has strengths but also has some weaknesses like connection and dependency. The system uses an external API and this can cause problems. Firstly the system cannot work offline as it needs internet to call the external API and this can be seen as a weakness. Secondly what happens if the external API is down for some reasons? this means that we cannot use the current system too. These weaknesses can be solved by building a full annotation system which isn’t dependent of any available external annotation API.

In the future, we think we can improve algorithms to handle grammar problems and disambiguation. This algorithms should use language dictionaries to be able to transform terms without meaning (short forms) to something understandable for better matching to concepts. For example when having an abbreviated term, there should be a dictionary to look up this term and return the full meaning. This will further help to reduce the problem of grammar mentioned earlier.

## Acknowledgments

Authors thank ICTLab for their support.

## References

1. P. Oliveira, J. Rocha. Semantic annotation tools survey. In: 2013 IEEE Symposium on Computational Intelligence and Data Mining (CIDM). 2013. p. 301–7.

2. Liao Y, Lezoche M, Panetto H, Boudjlida N. Why, Where and How to use Semantic Annotation for Systems Interoperability. In: 1st UNITE Doctoral Symposium. Bucarest, Romania; 2011. p. 71–8. https://hal.archives-ouvertes.fr/hal-00597903.

3. Cooper L, Meier A, Laporte MA, Elser JL, Mungall C, Sinn BT, et al. The Planteome database: An integrated resource for reference ontologies, plant genomics and phenomics. Nucleic Acids Res. 2018;46.

4. Jovanović J, Bagheri E. Semantic annotation in biomedicine: the current landscape. J Biomed Semant. 2017;8:44.

5. Jonquet C, Shah NH, Musen M a. The open biomedical annotator. Summit Transl Bioinforma. 2009;2009:56–60.

6. Noy NF, Shah NH, Whetzel PL, Dai B, Dorf M, Griffith N, et al. BioPortal: ontologies and integrated data resources at the click of a mouse. Nucleic Acids Res. 2009;37 Web Server issue:W170–173.

7. Jonquet C, Dzalé-Yeumo E, Arnaud E, Larmande P. AgroPortal: a proposition for ontology-based services in the agronomic domain. In: IN-OVIVE’15: 3{è}me atelier INt{é}gration de sources/masses de donn{é}es h{é}t{é}rog{è}nes et Ontologies, dans le domaine des sciences du VIVant et de l’Environnement. 2015.

